# The Effects of Learnability and Reward Responsiveness on Reward Processing

**DOI:** 10.64898/2026.04.13.718323

**Authors:** Abigail Oloriz, Olave E. Krigolson, Cameron D. Hassall

## Abstract

For methodological reasons, reward processing is commonly studied using random feedback and unlearnable tasks. It remains unclear whether task learnability influences reward-related brain activity, and whether this effect depends on individual differences such as reward responsiveness. We addressed this question by administering a behavioural activation system (BAS) scale before recording electroencephalography (EEG) while participants completed learnable and unlearnable versions of the “doors” task, a standard two-choice paradigm. Despite matched outcome likelihoods across conditions, participants reported greater motivation, enjoyment, and perceived performance in the learnable task. Contrary to our predictions, the amplitude of the reward positivity (RewP) – a frontocentral ERP index of reward processing – did not depend on task learnability and reward responsiveness. However, learnability and reward responsiveness effects became apparent when the analysis was restricted to high performers. Within this subgroup, participants low in reward responsiveness showed an enhanced RewP when the task was learnable. These findings suggest that contextual factors such as task learnability can interact with individual differences, informing ongoing efforts to identify the RewP as a biomarker of disordered reward processing.

---

Adaptive behaviour depends on learning which actions lead to reward. Early learning theory described this with Thorndike’s law of effect, which states that behaviours that lead to rewarding outcomes are more likely to be repeated, whereas unrewarded behaviours decrease in frequency (Thorndike, 1898). More recent computational models propose that learning is driven by reward prediction errors, or the difference between actual and expected outcomes (Rescorla & Wagner, 1972; Schultz et al., 1997; Sutton & Barto, 2018). When outcomes are better than expected, prediction errors strengthen action-outcome associations, and when they are worse than expected, those associations weaken. Taken together, these perspectives suggest that reward processing supports learning by updating expectations that can then be used to optimize future behaviour.

In humans, reward processing can be measured non-invasively using electroencephalography (EEG). For example, one component of the event-related potential (ERP) is the reward positivity (RewP^1^), a frontocentral signal that emerges approximately 240-340 ms following feedback and is reliably larger for rewards relative to non-rewards (Proudfit, 2015). Converging evidence indicates that the RewP reflects activity within anterior cingulate cortex (ACC) and represents the cortical expression of a dopaminergic prediction error (Holroyd & Coles, 2002; Proudfit, 2015; Sambrook & Goslin, 2015; Walsh & Anderson, 2012). Consistent with this view, RewP amplitude has been shown to vary as a function of reward expectancy within experimental tasks (Krigolson, 2018; Sambrook & Goslin, 2015; Walsh & Anderson, 2012). As a result, the RewP has become a widely used index of feedback evaluation, and has been increasingly used in research examining individual differences in reward sensitivity and affective disorders such as depression (Nelson et al., 2016; Proudfit, 2015).

One of the most widely used paradigms to elicit the RewP is the “doors” task, a two-alternative forced choice task in which participants select between two identical doors on each trial and then receive feedback indicating either a monetary gain (i.e., a green arrow pointing up) or loss (i.e., a red arrow pointing down). The doors task has been widely adopted to examine contextual effects on reward processing and individual differences in reward sensitivity (Hajcak et al., 2006; Holroyd & Coles, 2002; Proudfit, 2015; Sambrook & Goslin, 2015). For example, perceived agency over outcomes modulates the RewP, eliciting larger amplitudes when feedback is contingent on behaviour and reduced amplitudes when outcomes occur independently of choice (Hassall et al., 2019). The overall value of the task also shapes reward processing – the RewP is enhanced during low-value tasks and reduced during high-value tasks (Hassall et al., 2022). Similarly, individuals who self-report greater reward responsiveness show larger RewP amplitudes (Bress et al., 2012), suggesting that the signal is sensitive to motivational factors.

Finally, individual differences in affective functioning influence reward sensitivity, with blunted RewP responses observed in depression and altered feedback processing reported across anxiety-related and obsessive-compulsive symptoms (Hajcak et al., 2006; Nelson et al., 2016; Proudfit, 2015). These findings suggest that the RewP reflects an integrated evaluation of reward that emerges from a combination of individual traits and contextual factors such as motivation, agency, and task value, rather than a uniform neural response to reward.

A defining feature of the doors task is that outcomes are unrelated to performance – that is, feedback is random and unassociated with a participant’s actions. This is desirable methodologically because it means that gains and losses occur with equal probability, thus avoiding confounding the RewP with probability-sensitive components such as the P300 (Krigolson, 2018). However, dissociating outcomes from actions also means that the widely used doors task is unlearnable. Consequently, and despite the theory that the RewP reflects a learning signal (Holroyd & Coles, 2002), relatively little work has directly examined whether reward processing depends on task learnability, defined as the extent to which a task allows participants to acquire action-outcome associations. The reinforcement learning theory of the RewP proposes that feedback generates a neural teaching signal when outcomes deviate from expected action values, allowing action values to be updated over time (Holroyd & Coles, 2002; Sambrook & Goslin, 2015). However, reliable reward predictions cannot form when outcomes are random and independent of choice. The consequences of this have been shown in previous work. In a task in which participants could infer reward probabilities by learning a rule, Bellebaum and Daum (2008) found that the RewP became sensitive to outcome expectancy only after participants had learned the task contingencies, with larger responses to unlikely negative outcomes in learners but not in non-learners. Similarly, in a perceptual categorization task requiring feedback-based learning, the feedback-locked response diminished as predictions became reliable, an effect absent in participants who failed to learn the task (Krigolson et al., 2009). Consistent with this pattern, Hassall et al. (2022) reported that individuals who were more successful in learning reward contingencies tended to show larger RewP effects. Together, these findings suggest that reward-related neural signals are closely tied to the ability to learn task contingencies, raising the possibility that the RewP depends on task learnability.

In addition to informing theories about the RewP, determining the effect of task learnability on reward processing may also be clinically relevant. Simple gambling paradigms such as the doors task have been used to discover reward-processing differences across a range of internalizing conditions. For example, blunted RewP responses are consistently associated with depression (Hajcak et al., 2003, 2006; Nelson et al., 2016; Proudfit, 2015). As a result of this work, the RewP has been proposed as a neural marker of disrupted reward processing.

Behaviourally, these disorders are often characterized using reinforcement sensitivity theory and operationalized using the behavioural inhibition system and behavioural activation system (BIS/BAS) scale, which includes a subscale for reward responsiveness (Carver & White, 1994; Gray & McNaughton, 2003). The RewP scales with BAS and its subscales, further supporting its identification as a biomarker of reward disorders (Bress & Hajcak, 2013; Lange et al., 2012; Pegg et al., 2021). However, because this work relies on the standard unlearnable doors task, the role of learnability in disordered reward processing remains unclear. Motivating the present work, we speculate that learnable tasks with clear contingencies may preferentially engage BAS-related processes compared to unlearnable tasks. In other words, BAS-related RewP effects may be driven by perceived task learnability, instead of (or in in addition to) a general reduction in reward responsiveness.

Research on intrinsic motivation provides further rationale for examining task learnability in reward processing. In environments where outcomes are informative and progress can be made, individuals can update their expectations and experience a sense of mastery, which promotes sustained engagement. Previous literature has shown that task engagement increases when individuals experience learning progress, with greater progress associated with flow, a state of deep task engagement associated with feelings of control and enjoyment, and improved task-related cognitive control (Bakker et al., 2011; Brändle et al., 2025; Csikszentmihalyi, 1990; Lu et al., 2025). Likewise, large-scale behavioural analyses of video game play show that people report the greatest enjoyment when tasks are neither trivial nor impossible but instead allow meaningful learning and improvement over time (Brändle et al., 2025). In contrast, when outcomes appear random or unrelated to behaviour, opportunities for learning diminish, which can reduce engagement and motivation. However, contrasting learnable and unlearnable tasks is challenging due to methodological concerns. As described earlier, the RewP overlaps with components sensitive to outcome frequency (Krigolson, 2018), and learning naturally alters these probabilities over time, confounding whether effects reflect contingency learning or simple statistics.

The present study examined whether reward-related neural responses differ between learnable and unlearnable tasks. EEG was recorded while participants completed two versions of the doors task. In learnable blocks, one door was associated with a higher probability of reward than the other, allowing the opportunity for participants to learn which option was more rewarding over time. In unlearnable blocks, outcomes were random and unrelated to participants’ choices. Surveys were used to measure individual differences in reward responsiveness as well as task engagement. This allowed us to directly test the effects of task learnability and reward responsiveness on both task engagement and the RewP.

Based on prior work suggesting that learnable tasks promote motivation and engagement, we hypothesized that participants would report greater motivation and engagement during learnable blocks compared to unlearnable blocks (Brändle et al., 2025; Lu et al., 2025). Linking this with prior ERP studies showing learning-related changes in RewP amplitude, we further hypothesized that reward-related neural responses would be enhanced for learnable blocks (Bellebaum & Daum, 2008; Hassall et al., 2022; Krigolson, 2018). Finally, given the evidence that individuals high in reward responsiveness show robust neural responses regardless of task context (Bress et al., 2012; Pegg et al., 2021), we hypothesized that the effect of learnability on the RewP would depend on the reward responsiveness BAS subscale. Specifically, we predicted that participants low in reward responsiveness would show a larger learnability effect on the RewP (learnable RewP minus unlearnable RewP), whereas participants high in reward responsiveness would show smaller differences between task conditions.

## Methods

### Participants

We recruited 40 undergraduate students from MacEwan University (13 male, 4 left-handed, 1 ambidextrous, *M_age_ =* 21.58 years, *SD_age_* = 5.01). Participants were screened to ensure they had no known neurological impairments and normal or corrected-to-normal vision. Six participants had coarse or curly hair, requiring more electrolytic gel to lower electrode impedances. All six were included in the study.

The sample size was determined based on previous work examining the RewP. Williams et al. (2021) reported a large effect size for the RewP (Cohen’s *d* = 0.9). A power analysis conducted in G*Power (Version 3.1.97) indicated that a sample size of *N* = 15 would be sufficient to achieve statistical power of .95 at an alpha value of .05. To ensure adequate power to detect smaller effects and to account for potential data exclusions because of EEG artifacts, we aimed for a target sample of *N* = 40 participants.

Two participants were excluded from the final dataset due to the presence of excessive EEG artifacts for a final *N* of 38. These participants would have required more than three electrodes to be interpolated according to our artifact detection procedure, discussed later. Participants were recruited from the MacEwan University psychology participant pool and received 3% course credit per hour in compensation. In addition, all participants received a performance-dependent monetary bonus (*M_bonus_ =* CA$4.49, *SD_bonus_* = CA$2.05). The study was approved by the MacEwan University Research Ethics Board, and all participants provided written informed consent prior to participation.

### Apparatus and Procedure

Participants were seated approximately 700mm from an LCD display (ASUS PG27AQDM; 588 x 321 mm; 60 Hz refresh rate; 2560 by 1440 pixels). Visual stimuli were presented using Psychtoolbox (Brainard, 1997; Pelli, 1997) implemented in MATLAB (Version R2024a, The MathWorks Inc.). Participants were given the option of listening to lyric-free music during the task, which was delivered through a USB audio interface (Behringer XENYX 802S) connected to studio monitors (ADAM Audio D3V). Participants were instructed both verbally and in writing to minimize head movements and eye movements during the task to reduce EEG artifacts. During the EEG setup, participants completed the BIS/BAS scale as developed by Carver and White (1994).

Participants then completed the doors task, a computer-based two-choice guessing game (Proudfit, 2015). On each trial, prior to stimulus onset, a white fixation dot (0.15° of visual angle wide) was presented on a grey background for 400-600 ms. Participants were instructed to maintain fixation on the central dot and minimize eye and head movements during the task. Two doors, each 2° of visual angle wide, were then presented on the screen. The door centers were positioned 1.5° to the left and right of the fixation dot (distance between doors: 1° of visual angle). Using a standard keyboard, participants selected the left door by pressing the ‘f’ key with their left index finger or the right door by pressing the ‘j’ key with their right index finger. Participants were given a 2-second time limit to respond, otherwise they would see the message “TOO SLOW”. Following the response and prior to visual feedback, the doors disappeared and another grey background with white fixation dot was presented for 400**–**600 ms. Visual feedback indicating the trial outcome was then presented for 1000 ms as either an upward green arrow (reward) or downward red arrow (no reward), each 1° of visual angle wide. If the selected door contained a reward, the participant gained $0.05; otherwise, they lost $0.02. Participants were informed that their total earnings would be paid out at the end of the experiment. The task would then loop back to the next trial until the end of the block (Fig. 1a). Self-paced rest breaks were provided between each block.

**Figure 1.**
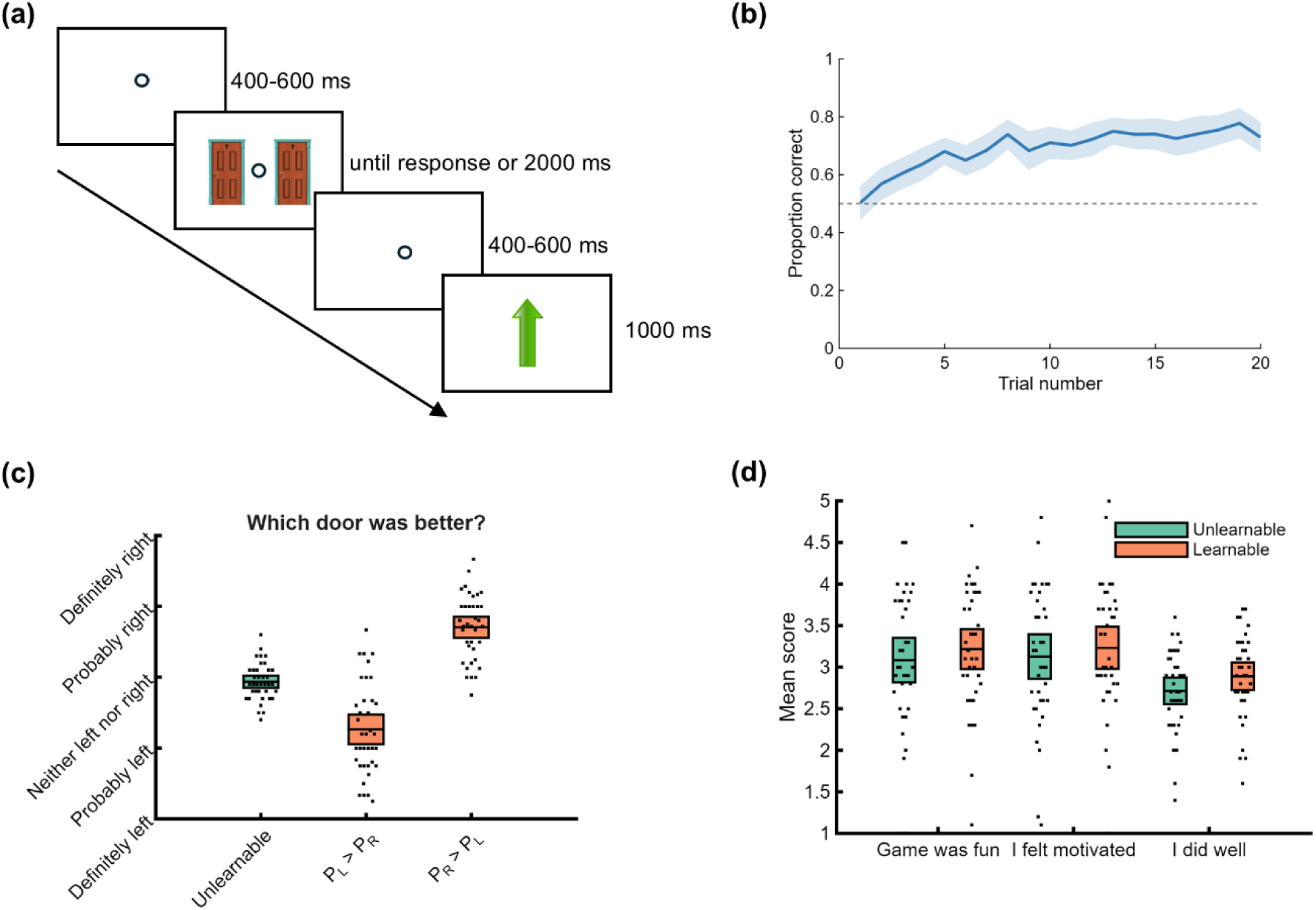
Task design and behavioural results. (a) Example trial sequence in the doors task. Participants selected between two doors, followed by feedback indicating reward (green upward arrow, +$0.05) or non-reward (red down arrow, –$0.02). Learnable blocks had one door with 60% probability and one with 10% probability; unlearnable blocks shuffled these outcomes randomly. (b) Learning curve showing proportion of optimal door selections in learnable blocks. Performance in the learnable blocks increased above chance after initial trials. (c) Post-block ratings of door preference. Participants correctly identified the better door in learnable blocks. (d) Subjective engagement ratings (fun, motivation, perceived performance). All ratings were higher in learnable vs. unlearnable blocks. Shaded region and error bars represent 95% confidence intervals.

The task consisted of 20 blocks of 20 trials each. Unbeknownst to participants, there were two block types: learnable and unlearnable (10 of each). In learnable blocks, one door (left or right) was associated with a higher probability of reward than the other (0.6 versus 0.1). These reward probabilities have been used previously in a large-sample RewP study, and have been shown to yield roughly equal numbers of wins and losses over a 20-trial block (Williams et al., 2021). The reward probabilities were assigned randomly to the two doors at the start of each learnable block. In unlearnable blocks, outcomes corresponded to reward sequences generated in prior learnable blocks but presented in random order and not linked to participant actions. The task always began with a learnable block followed by an unlearnable block. Remaining blocks were presented in a pseudorandom order such that no more than two blocks of the same type occurred consecutively. Thus, the overall reward frequency was preserved between block types.

Following each block, participants completed a brief survey to assess contingency awareness and task engagement. Participants were prompted to indicate which door they believed led to better outcomes using a five-point response scale (“definitely left”, “probably left”, “neither left nor right”, “probably right”, “definitely right”). Participants also rated agreement with the statements “The game was fun”, “I felt motivated to play the game”, and “I did well in the game” using five-point Likert scales. Question order was randomized across blocks.

### Data Collection

On each trial, the experimental software recorded behavioural data including participant response (left or right), response time (time since appearance of doors), trial outcome (reward or non-reward), and block type (learnable, unlearnable). With a 32-channel electrode cap referenced to Fz, EEG was recorded using BrainVision Recorder (Version 1.23.0001, Brain Products GmbH). Additional electrodes were placed on the left and right mastoids for later referencing. Electrodes were applied using conductive gel to reduce impedances and ensure a high-quality recording. All electrode impedances were reduced to below 20 kΩ prior to starting the recording. The EEG was then sampled at 1000 Hz and amplified using a BrainAmp DC amplifier (Brain Products GmbH) with a 250 Hz anti-aliasing filter applied during acquisition.

### Data Analysis

#### Behavioural Analysis

Participant responses on the BIS/BAS scale were transcribed and each subscale totaled, including an aggregate BAS score. To generate a learning curve, we calculated – for each trial – the mean proportion of optimal door selections across all learnable blocks. Participants’ explicit knowledge of which door had a higher probability of reward was assessed using post-block survey responses to the question “Which door was better?” and was averaged separately for each participant and block type (unlearnable, learnable). Subjective engagement was assessed by collecting mean Likert-scale ratings of fun, motivation, and perceived performance for each participant and block type (unlearnable, learnable).

#### EEG Preprocessing

The EEG was analyzed in MATLAB (Version R2025b, The MathWorks Inc.) and confirmed with BrainVision Analyzer (Version 2.3.0, Brain Products GmbH). EEG data was downsampled to 250Hz, bandpass filtered (0.1-30Hz, 60Hz notch), then re-referenced to the average of the mastoid signals. Ocular artifacts were detected using a MATLAB toolbox called *iclabel*, which relies independent component analysis (ICA). The ICA was trained using the complete recording, excluding segments where the voltage surpassed 1000 μV or dipped below – 1000 μV. On average, *iclabel* identified 2.75 ocular components (*SD* = 0.95), which were removed from the EEG.

We then constructed 800 ms epochs around each feedback event (–200 to 600 ms). Baseline correction was applied from –200 to 0 ms. The data were subsequently examined for any lingering artifacts – that is, any epochs that exhibited a voltage change exceeding 10 μV per sample point or an overall change in voltage of more than 150 μV. Feedback locked epochs with such artifacts were rejected. Artifact rejection was done separately for each electrode, which were labelled as “noisy” if more than 10% of their epochs were rejected. Participants with more than three noisy electrodes were exclude from subsequent analysis (two participants in total).

Noisy electrodes were then interpolated for the remaining participants (*M* = 0.08 interpolated electrodes, *SD* = 0.27). For each remaining participant, we then constructed ERPs by averaging the EEG across the remaining artifact-free epochs for each outcome (win, loss) and condition (unlearnable, learnable). To quantify the RewP, we employed the difference wave method by subtracting the average loss ERP from the average win ERP for each participant and block type (unlearnable, learnable). The difference wave method is particularly effective for isolating the RewP as it reduces component overlap (Krigolson, 2018; Luck, 2014). We then analyzed the electrode and the time interval suggested by Sambrook and Goslin (2015): FCz from 240 to 340 ms after feedback. The RewP was identified as the mean voltage measured within this time frame for each participant and block type (unlearnable, learnable).

#### Statistics

##### Behavioural and RewP Analyses

Engagement ratings (fun, motivation, and perceived performance) were compared between conditions (unlearnable, learnable) using paired-samples *t*-tests. RewP amplitude was also compared between learnable and unlearnable conditions using paired-samples t-tests. To examine whether reward responsiveness moderated learnability effects, BAS reward responsiveness scores were correlated with the learnability-related RewP difference score (learnable minus unlearnable). The BIS/BAS scale comprises four subscales: one score for BIS and three for BAS (drive, reward responsiveness, and fun seeking). Previous literature suggests that all three BAS types as well as the aggregate BAS score correlate with reward processing. Here we focused on reward responsiveness as this subscale has been shown to be particularly clinically relevant (Taubitz et al., 2015). However, we repeated our statistical tests for each of the BIS/BAS subscales as well as the aggregate BAS score.

##### Exploratory Analyses

An exploratory multiple regression tested whether task performance moderated the relationship between reward responsiveness and the learnability-related RewP difference score. Task performance was indexed using win-stay probability, defined as the proportion of trials on which participants repeated the same choice following rewarded actions. This method of measuring performance has been used previously to measure performance in the doors task, and is particularly appealing here because it can be applied to both unlearnable and learnable trials (Bowyer et al., 2022). Participants were divided into tertiles based on their win-stay probability, such that individuals in the upper third of the distribution were classified as high performers and those in the lower third were classified as low performers. These groupings were used to visualize the effects of task performance and reward responsiveness on reward processing.

## Results

### Behavioural Results

To examine whether participants learned the reward contingencies in the learnable condition, accuracy across trials was plotted as a function of trial number (Fig. 1b). Performance increased across trials and remained above chance levels after the early portion of the block, indicating that participants were sensitive to the underlying reward structure. Furthermore, responses to the post-block question assessing which door was better indicated that participants correctly identified the higher-probability door in the learnable condition (Fig. 1c).

Participants reported greater task engagement in the learnable condition compared to the unlearnable condition. Specifically, ratings of fun were higher in the learnable block, *t*(36) = 2.68, *p* = .01, Cohen’s *d* = 0.44. Similarly, participants reported greater motivation in the learnable block, *t*(36) = 2.39, *p* = .02, Cohen’s *d* = 0.39, and higher perceived performance, *t*(36) = 2.44, *p* = .02, Cohen’s *d* = 0.40 (Fig. 1d). These results indicate modest but reliable differences in subjective engagement between task conditions.

### ERP Results

Across conditions, gain feedback elicited a more positive ERP response than loss feedback, confirming the presence of a typical RewP effect (Fig. 2). However, RewP amplitude did not differ between the learnable and unlearnable conditions, *t*(37) = 0.59, *p* = .56, Cohen’s *d* = 0.17. Waveform morphology and scalp topography were similar across conditions, consistent with the absence of a learnability effect on the RewP.

**Figure 2.**
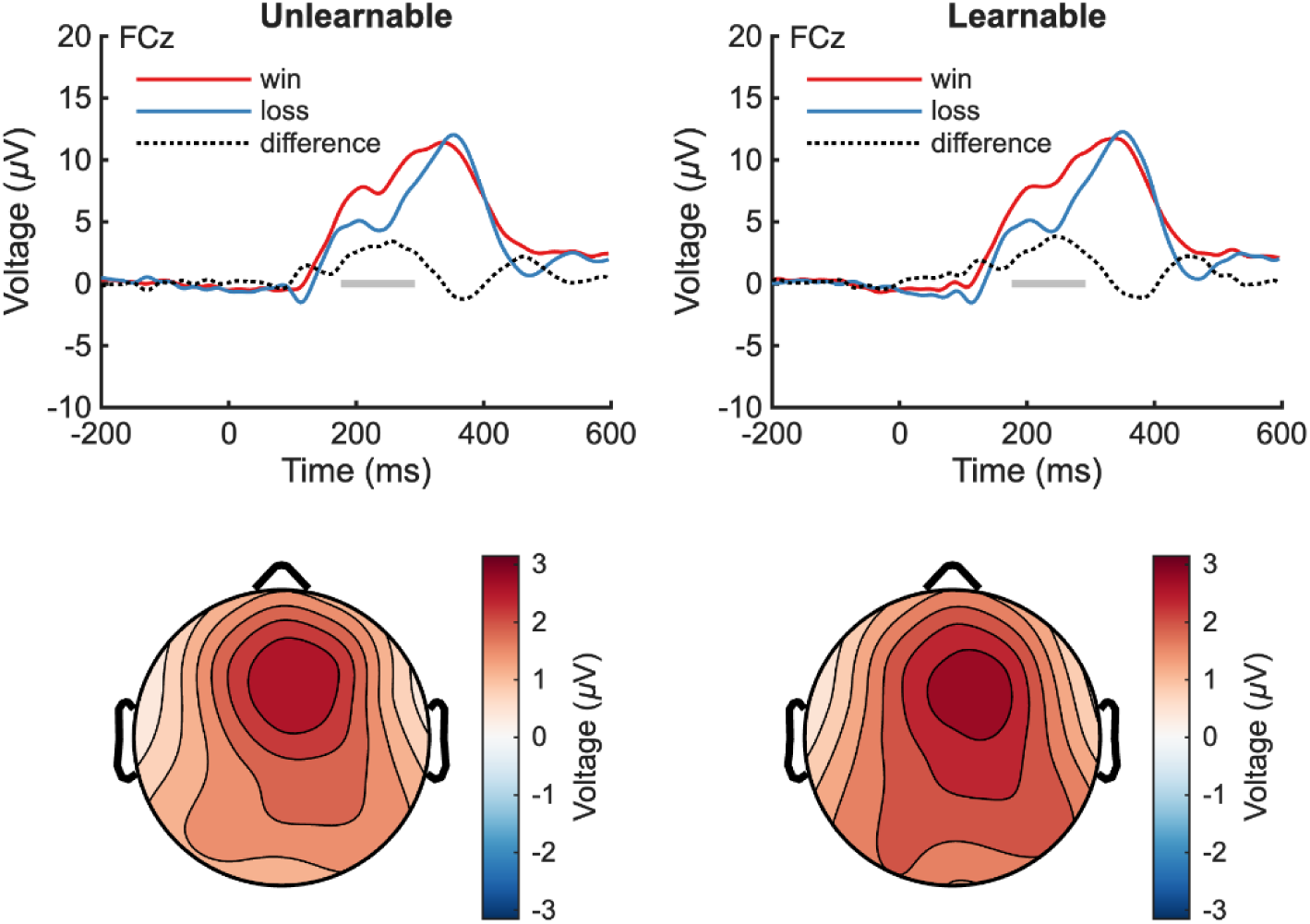
Grand average ERP waveforms and scalp topography for wins (rewards) and losses (non-rewards). Waveforms are shown at electrode FCz for the learnable and unlearnable conditions. Gain feedback elicited more positive ERPs than loss feedback in both conditions, confirming the presence of a typical RewP effect. Grey shading shows the analysis time window (240-340 ms post-feedback). Scalp topographies shows similar spatial distributions of the RewP effect across conditions.

To examine whether individual differences in reward sensitivity moderated the effect of learnability on reward processing, BAS reward responsiveness scores were correlated with the learnability-related △RewP (learnable – unlearnable). Contrary to predictions, this relationship was not significant, *r*(36) = 0.04, *p* = .81 (Fig. 3), indicating that participants with lower reward responsiveness did not show a larger learnability-related increase in RewP amplitude than participants with higher reward responsiveness. We also noted that △RewP (learnable – unlearnable) was uncorrelated with the different BAS subscales (see Supplementary Table 1). However, an exploratory analysis examining task performance suggested a more complex relationship between reward responsiveness, learnability, and the RewP (see below).

**Figure 3.**
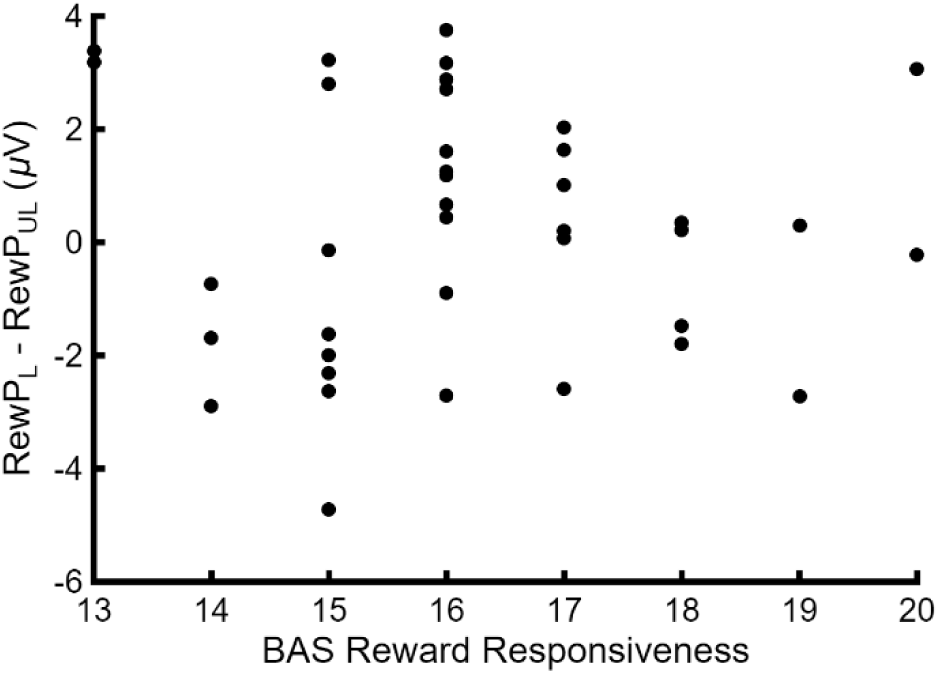
Scatterplot of BAS scores and the learnability-related RewP difference scores (learnable minus unlearnable). No significant correlation was observed, suggesting that reward responsiveness did not moderate the effect of learnability on RewP amplitude as predicted.

### Exploratory Results

Our exploratory analysis examined whether behavioural performance moderated the relationship between reward responsiveness and the learnability-related △RewP (learnable – unlearnable). A multiple regression model including reward responsiveness, task performance (win-stay probability), and their interaction significantly predicted variation in △RewP, *F*(3,34) = 3.35, *p* = .03 (adjusted *R*^2^ = 0.16). Neither reward responsiveness nor task performance independently predicted △RewP; however, there was a significant reward responsiveness by performance interaction, β = –0.36, *p* = .004 (Fig. 4), indicating that the relationship between reward responsiveness and the learnability-related RewP effect differed as a function of task performance. When we examined the overall BAS score and the other BIS/BAS subscales, this result was replicated for the aggregate BAS score and the fun seeking subscale, but not for drive or BIS (Supplementary Table 2).

**Figure 4.**
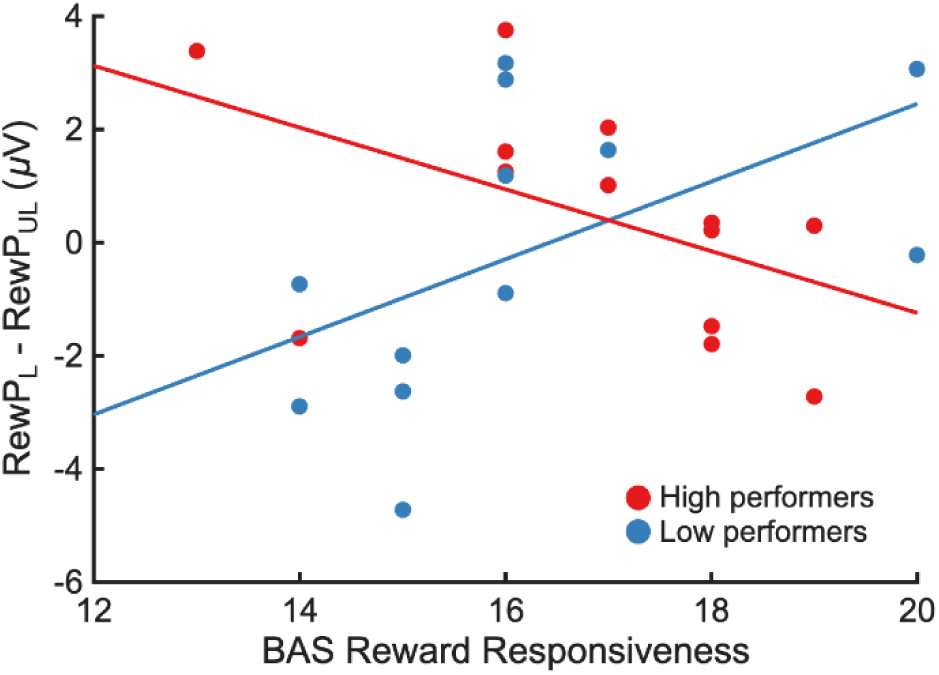
Reward responsiveness by task performance interaction on the learnability-related RewP effect (learnable minus unlearnable). The multiple regression model revealed a significant interaction. Higher-performing participants showed the predicted negative relationship between reward responsiveness and the learnability-related RewP effect, while lower-performing participants showed the opposite pattern.

## Discussion

The present study examined whether reward-related neural responses depend on task learnability, and whether this relationship is moderated by individual differences in reward responsiveness. Consistent with our first hypothesis, participants reported greater motivation and enjoyment during learnable trials compared to unlearnable trials, suggesting that environments that allow learning progress enhance task engagement. Contrary to our second and third hypotheses, however, reward processing did not depend on learnability or reward responsiveness.

The finding that participants reported greater fun, motivation, and perceived performance during learnable blocks compared to unlearnable blocks suggests that learnability enhances engagement in reward-based decision-making tasks. One possible explanation is that learnable tasks allow participants to experience progress toward a goal, which has been shown to improve intrinsic motivation (Gottlieb et al., 2013; Gottlieb & Oudeyer, 2018; Kaplan & Oudeyer, 2007).

Consistent with this interpretation, Lu et al. (2025) demonstrated that task engagement increases when individuals experience learning progress, with greater progress associated with higher levels of flow and reduced distractibility. Similarly, Brändle et al. (2025) provided evidence that people’s preference for moderately challenging environments depends on whether they experience learning progress within those environments, suggesting that enjoyment may arise from noticing improvement over time rather than maintaining the same perceived level of performance. This account aligns with the present finding that participants reported greater fun and motivation in the learnable condition, suggesting that opportunities to detect differences in door value across trials supported a more positive task experience. Despite matched outcome frequencies across block types, participants rated their performance more highly when outcomes were yoked to actions. This increase in perceived performance further suggests that participants were able to detect differences across conditions, possibly contributing to a greater sense of control over outcomes in learnable trials. Learnable environments may therefore enhance engagement not only by supporting learning progress but also by increasing perceived control over outcomes, which has been shown to enhance reward processing (Hassall et al., 2019; Mühlberger et al., 2017). Together, these findings suggest that tasks that do not support learning may be experienced as less engaging and motivating, even when reward outcomes themselves are held constant.

Our EEG analysis revealed strong evidence of reward processing in both of our conditions. The presence of a RewP in the unlearnable condition is unsurprising as previous research suggests that reward-related feedback responses such as the RewP are observed even when individuals are aware that the chance of reward is random (Mühlberger et al., 2017). Indeed, the standard doors task reliably elicits a RewP even though feedback is random, indicating that participants continue to differentiate gains from losses in the absence of learnable action-outcome contingencies (Bress & Hajcak, 2013; Lange et al., 2012; Pegg et al., 2021). Although previous work clearly predicts the presence of a RewP in the unlearnable condition, we further predicted that the RewP would be enhanced if participants perceived learning progress across trials. This hypothesis was not supported by our data, a result that is incongruous with previous work suggesting that reward-related neural signals are sensitive to learning and expectancy updating across individuals (Bellebaum & Daum, 2008; Hassall et al., 2022; Krigolson et al., 2009). One possible explanation for the absence of a learning-related RewP enhancement is that our learnability manipulation produced only small-to-moderate behavioural effects, suggesting that the extent to which participants experienced the task environments as meaningfully distinct was limited. Similarly, and contrary to our third hypothesis, individual differences in reward responsiveness did not moderate the effect of learnability on the RewP amplitude, as BAS reward responsiveness scores were not directly associated with differences in RewP amplitude between the learnable and unlearnable trials. This does not align with previous work showing a correlation between BAS and RewP scores (Bress & Hajcak, 2013; Lange et al., 2012; Pegg et al., 2021), though it’s worth pointing out that not all previous studies have observed this relationship (Balconi & Crivelli, 2010; Cooper et al., 2014).

Our exploratory analysis may provide additional explanation for these null effects. It suggests the relationship between learnability and reward processing depends on both reward responsiveness and task performance. Specifically, BAS interacted with win-stay behaviour, such that higher-performing participants showed the predicted effect of BAS and learnability on the RewP, whereas lower-performing participants showed an effect in the opposite direction. This pattern may have appeared because higher-performing participants were more likely to detect the difference in outcome probabilities between doors in the learnable condition, allowing reward signals to become more informative and more strongly reflected in the RewP. In contrast, lower-performing participants may not have detected this structure and may have instead responded more randomly. Together, these findings suggest that individual differences in strategy use and task engagement can influence how learnability interacts with reward responsiveness to shape neural responses to feedback. Further study will be needed to verify and explain these effects.

The present observations may have implications for the use of the RewP as a biomarker in clinical research. The RewP is widely used as an index of reward responsiveness and has been proposed as a neural marker of internalizing disorders such as depression (Chen et al., 2015; De Aguiar Neto & Rosa, 2019; Hajcak et al., 2003, 2006; Keren et al., 2018; Liang et al., 2025; Mackin et al., 2021; Mukherjee et al., 2020; Nelson et al., 2016; Nielson et al., 2021; Proudfit, 2015; Umemoto & Holroyd, 2017). The doors task is commonly used to measure the RewP in this work precisely because outcomes are random and unrelated to performance, which helps to isolate the neural response to rewards. However, our findings suggest that participants may differ in how they respond to task learnability. For some individuals, RewP amplitude could partly reflect task engagement due to perceived learnability rather than a pure measure of reward sensitivity. Task learnability may thus be an important contextual factor to consider when interpreting blunted RewP responses in clinical populations.

Several limitations should be considered when interpreting these findings. First, the reward responsiveness by performance interaction we observed emerged from an exploratory analysis and requires replication. In addition, an undergraduate sample may limit generalizability to broader populations with greater reward responsiveness variability. Next, small-to-moderate behavioural effects suggest that our learnability manipulation may have been too subtle for all participants to detect reliably. Finally, relatively short blocks of 20 trials may have limited opportunities for robust learning and strategy development. Future research with stronger manipulations, larger/diverse samples, and extended learning periods could further examine how task learnability shapes reward processing.

In summary, the present results suggest that task learnability subtly influences reward processing by interacting with reward responsiveness and strategy use, rather than producing large changes at the group level. These findings contribute to the growing understanding that reward processing depends on both individual factors and task context, with implications for the interpretation of clinical biomarkers of depression and other reward-related disorders.

## Author Note

This research was funded by a Natural Sciences and Engineering Research Council of Canada (NSERC) Discovery Grant to Cameron D. Hassall (RGPIN 2024-04848).

The authors have no conflict of interest.

## Author Contributions

**Abigail Oloriz:** Conceptualization; formal analysis; investigation; writing – original draft preparation; writing – review and editing. **Olave E. Krigolson:** Conceptualization; writing – review and editing. **Cameron D. Hassall:** Conceptualization; formal analysis; investigation; data curation; writing – review and editing; supervision.

## Data Availability Statement

EEG dataset is available at https://openneuro.org/datasets/ds007647/versions/1.0.0

Analysis scripts are available at https://github.com/chassall/differentdoors.

## Footnotes

1 Historically, the RewP has been referred to as the feedback-related negativity (FRN), the medial frontal negativity (MFN), the feedback error-related negativity (fERN), and the feedback negativity (FN). Accumulating evidence suggests that this signal is more accurately characterized as a reward-related positivity driven by gains rather than a negativity driven by losses, leading to its reconceptualization as the RewP (Proudfit, 2015).

## Notes

https://openneuro.org/datasets/ds007647/versions/1.0.0

### Competing Interest Statement

The authors have declared no competing interest.

### Summary of Updates

Minor edits to the text and main figures. Supplement is unchanged.

## References

1. Bakker, A. B., Oerlemans, W., Demerouti, E., Slot, B. B., & Ali, D. K. (2011). Flow and performance: A study among talented Dutch soccer players. Psychology of Sport and Exercise, 12(4), 442–450. 10.1016/j.psychsport.2011.02.003

2. Balconi, M., & Crivelli, D. (2010). FRN and P300 ERP effect modulation in response to feedback sensitivity: The contribution of punishment-reward system (BIS/BAS) and Behaviour Identification of action. Neuroscience Research, 66(2), 162–172. 10.1016/j.neures.2009.10.011

3. Bellebaum, C., & Daum, I. (2008). Learning-related changes in reward expectancy are reflected in the feedback-related negativity. European Journal of Neuroscience, 27(7), 1823–1835. 10.1111/j.1460-9568.2008.06138.x

4. Bowyer, C. B., Brush, C. J., Patrick, C. J., & Hajcak, G. (2022). Effort and Appetitive Responding in Depression: Examining Deficits in Motivational and Consummatory Stages of Reward Processing Using the Effort-Doors Task. Biological Psychiatry Global Open Science, 3(4), 1073–1082. 10.1016/j.bpsgos.2022.08.002

5. Brainard, D. H. (1997). The Psychophysics Toolbox. Spatial Vision, 10(4), 433–436. 10.1163/156856897X00357

6. Brändle, F., Wu, C. M., & Schulz, E. (2025). Leveling up fun: Learning progress, expectations, and success influence enjoyment in video games. Scientific Reports, 15(1), 34153. 10.1038/s41598-025-14628-2

7. Bress, J. N., & Hajcak, G. (2013). Self-report and behavioral measures of reward sensitivity predict the feedback negativity. Psychophysiology, 50(7), 610–616. 10.1111/psyp.12053

8. Bress, J. N., Smith, E., Foti, D., Klein, D. N., & Hajcak, G. (2012). Neural response to reward and depressive symptoms in late childhood to early adolescence. Biological Psychology, 89(1), 156–162. 10.1016/j.biopsycho.2011.10.004

9. Carver, C. S., & White, T. L. (1994). Behavioral Inhibition, Behavioral Activation, and Affective Responses to Impending Reward and Punishment: The BIS/BAS Scales. Journal of Personality and Social Psychology, 67(2), 319–333. 10.1037/0022-3514.67.2.319

10. Chen, C., Takahashi, T., Nakagawa, S., Inoue, T., & Kusumi, I. (2015). Reinforcement learning in depression: A review of computational research. Neuroscience & Biobehavioral Reviews, 55, 247–267. 10.1016/j.neubiorev.2015.05.005

11. Cooper, A. J., Duke, É., Pickering, A. D., & Smillie, L. D. (2014). Individual differences in reward prediction error: Contrasting relations between feedback-related negativity and trait measures of reward sensitivity, impulsivity and extraversion. Frontiers in Human Neuroscience, 8. 10.3389/fnhum.2014.00248

12. Csikszentmihalyi, M. (1990). Flow: The psychology of optimal experience. New York: Harper and Row.

13. De Aguiar Neto, F. S., & Rosa, J. L. G. (2019). Depression biomarkers using non-invasive EEG: A review. Neuroscience & Biobehavioral Reviews, 105, 83–93. 10.1016/j.neubiorev.2019.07.021

14. Gottlieb, J., & Oudeyer, P.-Y. (2018). Towards a neuroscience of active sampling and curiosity. Nature Reviews Neuroscience, 19(12), 758. https://doi.org/10/gfgzr9

15. Gottlieb, J., Oudeyer, P.-Y., Lopes, M., & Baranes, A. (2013). Information-seeking, curiosity, and attention: Computational and neural mechanisms. Trends in Cognitive Sciences, 17(11), 585–593. 10.1016/j.tics.2013.09.001

16. Gray, J. A., & McNaughton, N. (2003). The Neuropsychology of Anxiety: An enquiry into the function of the septo-hippocampal system (2nd ed.). Oxford University Press. 10.1093/acprof:oso/9780198522713.001.0001

17. Hajcak, G., McDonald, N., & Simons, R. F. (2003). Anxiety and error-related brain activity. Biological Psychology, 64(1–2), 77–90. 10.1016/S0301-0511(03)00103-0

18. Hajcak, G., Moser, J. S., Holroyd, C. B., & Simons, R. F. (2006). The feedback-related negativity reflects the binary evaluation of good versus bad outcomes. Biological Psychology, 71(2), 148–154. 10.1016/j.biopsycho.2005.04.001

19. Hassall, C. D., Hajcak, G., & Krigolson, O. E. (2019). The importance of agency in human reward processing. *Cognitive, Affective*, & Behavioral Neuroscience, 19(6), 1458–1466. 10.3758/s13415-019-00730-2

20. Hassall, C. D., Hunt, L. T., & Holroyd, C. B. (2022). Task-level value affects trial-level reward processing. NeuroImage, 260, 119456. 10.1016/j.neuroimage.2022.119456

21. Holroyd, C. B., & Coles, M. G. H. (2002). The neural basis of human error processing: Reinforcement learning, dopamine, and the error-related negativity. Psychological Review, 109(4), 679–709. 10.1037/0033-295X.109.4.679

22. Kaplan, F., & Oudeyer, P.-Y. (2007). In search of the neural circuits of intrinsic motivation. Frontiers in Neuroscience, 1. 10.3389/neuro.01.1.1.017.2007

23. Keren, H., O’Callaghan, G., Vidal-Ribas, P., Buzzell, G. A., Brotman, M. A., Leibenluft, E., Pan, P. M., Meffert, L., Kaiser, A., Wolke, S., Pine, D. S., & Stringaris, A. (2018). Reward Processing in Depression: A Conceptual and Meta-Analytic Review Across fMRI and EEG Studies. American Journal of Psychiatry, 175(11), 1111–1120. 10.1176/appi.ajp.2018.17101124

24. Krigolson, O. E. (2018). Event-related brain potentials and the study of reward processing: Methodological considerations. International Journal of Psychophysiology, 132, 175–183. 10.1016/j.ijpsycho.2017.11.007

25. Krigolson, O. E., Pierce, L. J., Holroyd, C. B., & Tanaka, J. W. (2009). Learning to Become an Expert: Reinforcement Learning and the Acquisition of Perceptual Expertise. Journal of Cognitive Neuroscience, 21(9), 1833–1840. 10.1162/jocn.2009.21128

26. Lange, S., Leue, A., & Beauducel, A. (2012). Behavioral approach and reward processing: Results on feedback-related negativity and P3 component. Biological Psychology, 89(2), 416–425. 10.1016/j.biopsycho.2011.12.004

27. Levinson, A. R., Speed, B. C., Infantolino, Z. P., & Hajcak, G. (2017). Reliability of the electrocortical response to gains and losses in the doors task. Psychophysiology, 54(4), 601–607. 10.1111/psyp.12813

28. Liang, Y., An, P., Liu, G., & Zheng, Y. (2025). Blunted perceived control effect on reward anticipation in trait anxiety. Journal of Affective Disorders, 388, 119559. 10.1016/j.jad.2025.119559

29. Lu, H., Van Der Linden, D., & Bakker, A. B. (2025). The neuroscientific basis of flow: Learning progress guides task engagement and cognitive control. NeuroImage, 308, 121076. 10.1016/j.neuroimage.2025.121076

30. Luck, S. J. (2014). An Introduction to the Event-Related Potential Technique (2nd ed.). MIT Press.

31. Mackin, D. M., Nelson, B. D., & Klein, D. N. (2021). Reward processing and depression: Current findings and future directions. In The Neuroscience of Depression (pp. 425–433). Elsevier. 10.1016/B978-0-12-817935-2.00051-9

32. Mühlberger, C., Angus, D. J., Jonas, E., Harmon-Jones, C., & Harmon-Jones, E. (2017). Perceived control increases the reward positivity and stimulus preceding negativity. Psychophysiology, 54(2), 310–322. 10.1111/psyp.12786

33. Mukherjee, D., Lee, S., Kazinka, R., D. Satterthwaite, T., & Kable, J. W. (2020). Multiple Facets of Value-Based Decision Making in Major Depressive Disorder. Scientific Reports, 10(1), 3415. 10.1038/s41598-020-60230-z

34. Nelson, B. D., Perlman, G., Klein, D. N., Kotov, R., & Hajcak, G. (2016). Blunted Neural Response to Rewards as a Prospective Predictor of the Development of Depression in Adolescent Girls. American Journal of Psychiatry, 173(12), 1223–1230. 10.1176/appi.ajp.2016.15121524

35. Nielson, D. M., Keren, H., O’Callaghan, G., Jackson, S. M., Douka, I., Vidal-Ribas, P., Pornpattananangkul, N., Camp, C. C., Gorham, L. S., Wei, C., Kirwan, S., Zheng, C. Y., & Stringaris, A. (2021). Great Expectations: A Critical Review of and Suggestions for the Study of Reward Processing as a Cause and Predictor of Depression. Biological Psychiatry, 89(2), 134–143. 10.1016/j.biopsych.2020.06.012

36. Pegg, S., Jeong, H. J., Foti, D., & Kujawa, A. (2021). Differentiating Stages of Reward Responsiveness: Neurophysiological Measures and Associations with Facets of the Behavioral Activation System. Psychophysiology, 58(4), e13764. 10.1111/psyp.13764

37. Pelli, D. G. (1997). The VideoToolbox software for visual psychophysics: Transforming numbers into movies. Spatial Vision, 10(4), 437–442. 10.1163/156856897X00366

38. Proudfit, G. H. (2015). The reward positivity: From basic research on reward to a biomarker for depression. Psychophysiology, 52(4), 449–459. 10.1111/psyp.12370

39. Rescorla, R. A., & Wagner, A. R. (1972). A theory of Pavlovian conditioning: Variations in the effectiveness of reinforcement and nonreinforcement. In A. H. Black & W. F. Prokasy (Eds.), Classical conditioning II: Current research and theory (pp. 64–99). Appleton-Century-Crofts.

40. Sambrook, T. D., & Goslin, J. (2015). A neural reward prediction error revealed by a meta-analysis of ERPs using great grand averages. Psychological Bulletin, 141(1), 213–235. 10.1037/bul0000006

41. Schultz, W., Dayan, P., & Montague, P. R. (1997). A Neural Substrate of Prediction and Reward. Science, 275(5306), 1593–1599. 10.1126/science.275.5306.1593

42. Sutton, R. S., & Barto, A. G. (2018). Reinforcement learning: An introduction (Second edition). The MIT Press.

43. Taubitz, L. E., Pedersen, W. S., & Larson, C. L. (2015). BAS Reward Responsiveness: A unique predictor of positive psychological functioning. Personality and Individual Differences, 80, 107–112. 10.1016/j.paid.2015.02.029

44. Thorndike, E. L. (1898). Animal intelligence: An experimental study of the associative processes in animals. The Psychological Review: Monograph Supplements, 2(4), i–109. 10.1037/h0092987

45. Umemoto, A., & Holroyd, C. B. (2017). Neural mechanisms of reward processing associated with depression-related personality traits. Clinical Neurophysiology, 128(7), 1184–1196. 10.1016/j.clinph.2017.03.049

46. Walsh, M. M., & Anderson, J. R. (2012). Learning from experience: Event-related potential correlates of reward processing, neural adaptation, and behavioral choice. Neuroscience & Biobehavioral Reviews, 36(8), 1870–1884. 10.1016/j.neubiorev.2012.05.008

47. Williams, C. C., Ferguson, T. D., Hassall, C. D., Abimbola, W., & Krigolson, O. E. (2021). The ERP, frequency, and time–frequency correlates of feedback processing: Insights from a large sample study. Psychophysiology, 58(2), e13722. 10.1111/psyp.13722

