## Supplement for "The Effects of Learnability and Reward Responsiveness on Reward Processing"

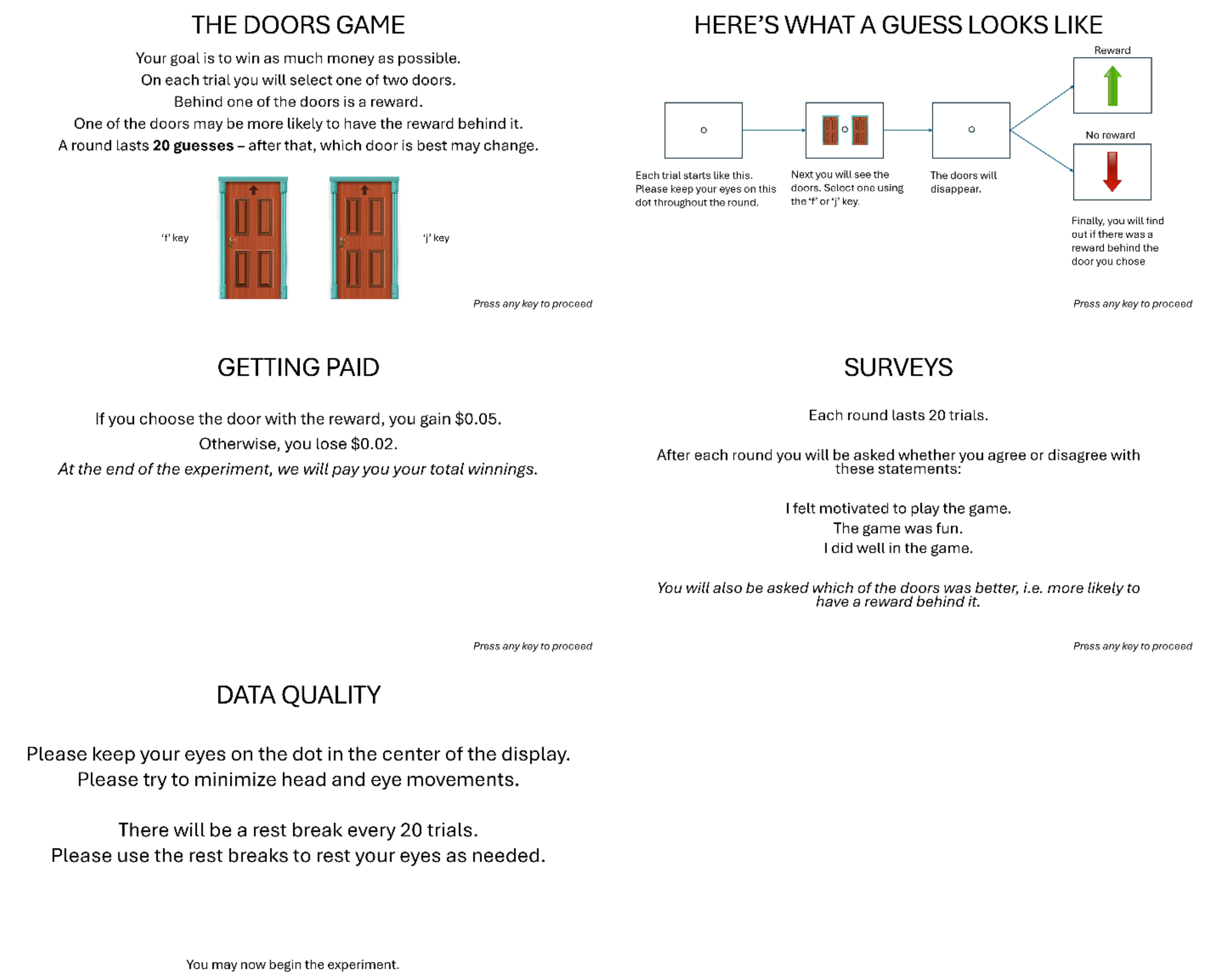


*Supplementary Figure 1*. Task instructions.

**Supplementary Table 1**

*Correlations between ΔRewP (learnable – unlearnable) and BIS/BAS scores*

| Subscale | *r* | *p* |
|---|---|---|
| BAS | 0.11 | .51 |
| BAS Drive | -0.02 | .92 |
| BAS Fun Seeking | 0.20 | .24 |
| BAS Reward Responsiveness | 0.04 | .81 |
| BIS | 0.22 | .18 |

**Supplementary Table 2**

*Multiple regression results for each BIS/BAS score*

| Subscale | *F* | *p* | Adjusted $R^2$ | Interaction β | *p* |
|---|---|---|---|---|---|
| BAS | 3.78 | .02 | 0.18 | -0.36 | .003 |
| BAS Drive | 0.82 | .49 | 0.01 | -0.21 | .18 |
| BAS Fun Seeking | 4.85 | .006 | 0.24 | -0.41 | .001 |
| BAS Reward Responsiveness | 3.35 | .03 | 0.16 | -0.36 | .004 |
| BIS | 0.88 | .46 | 0.01 | 0.15 | .42 |
